## Supplemental information for "Deep mutational scanning quantifies DNA binding and predicts clinical outcomes of PAX6 variants"

### SUPPLEMENTARY MATERIALS

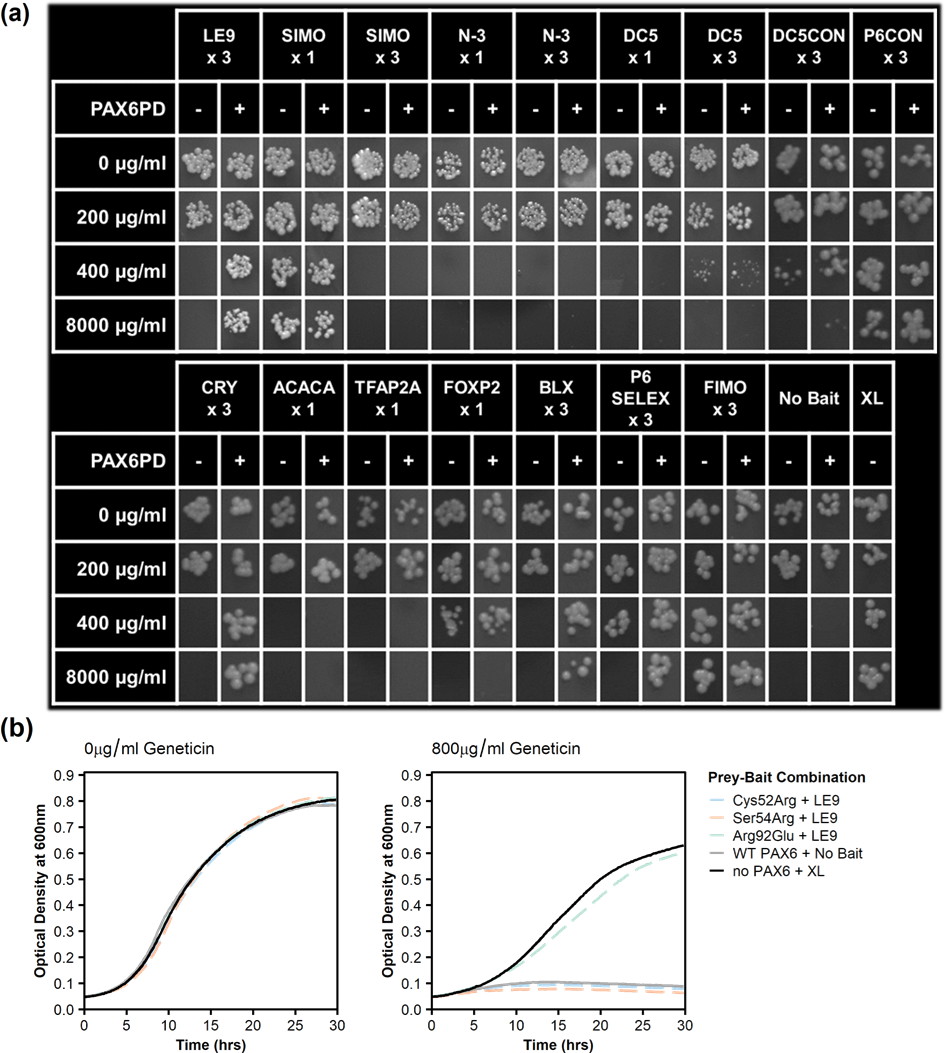

##### Fig S1. Bait-prey screening and characterised variant validation

(a) Spot assays used to screen PAX6:bait combinations. Single (x1) or three tandem repeats (x3) of PAX6 bait sequences were trialled alongside a strain lacking a bait sequence (No Bait), and synthetic minimal yeast promoter (XL) capable of independently driving expression of the Geneticin resistance gene. Each PAX6 bait-specific strain was screened with (+) and without (-) PAX6 expression over a range of Geneticin concentrations and serial dilutions summarised in each tile. (b) Liquid culture validation of the LE9 x3 bait strain using characterised PAX6 variants. Three PAX6 variants with varying levels of reported disruption to DNA-binding^1^ are represented as dashed lines. Log growth grate was measured by optical density.

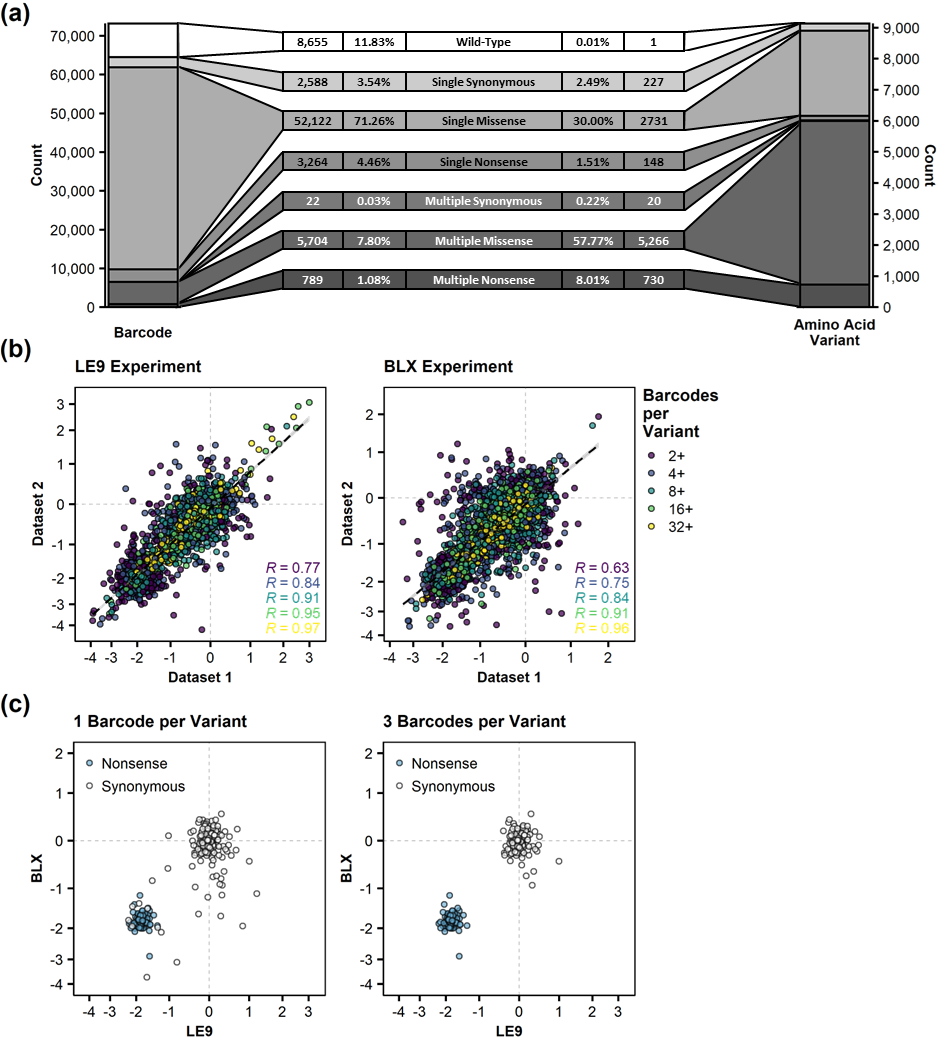

##### Fig S2. Variant library composition and barcode

(a) Library composition following saturation mutagenesis broken down by unique barcodes (left) and unique variants (right). The number of each variant by type and corresponding percentage of the total of each are shown. (b) Internal correlation between bins with increasing number of barcodes per variant and corresponding Pearson correlations for LE9 (left) and BLX (right). (c) Comparison of variant fitness scores of synonymous and nonsense variants in the LE9 and BLX assays, filtered by number of barcodes per variant (one, left versus three, right).

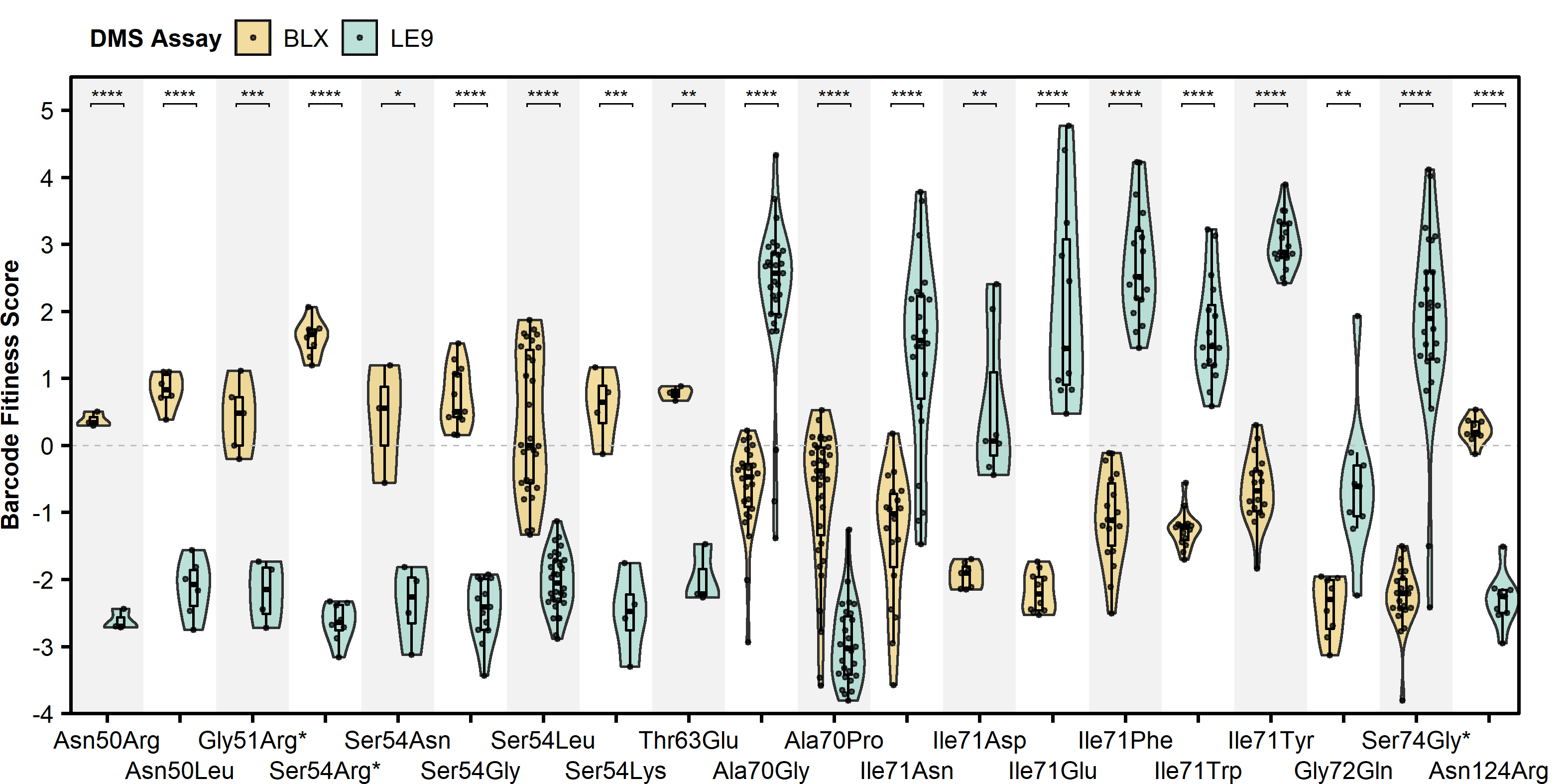

##### Fig S3. Variants with inverse changes in DNA-binding

The top 20 variants with the highest shifts in variant fitness score between the LE9 and BLX assay are shown. Black dots represent individual barcodes for each variant. Asterisks (*) on the x-axis denote pathogenic variants identified in human patients. All p-values are calculated using Wilcoxon test and adjusted using FDR.

| **Name** | **Sequence** | **Source** |
| --- | --- | --- |
| LE9 x 3 | ATGAGAGATCTTTCCGCTCATTGCCCATTCAAATACAATTGTAGATCATGAGAGATCTTTCCGCTCATTGCCCATTCAAATACAATTGTAGATCATGAGAGATCTTTCCGCTCATTGCCCATTCAAATACAATTGTAGATC | Aota et al.^2^ |
| SIMO x 1 | GTGTCAATGCCTGAAGTGATACGCTCTGA | Bhatia et al.^3^ |
| SIMO x 3 | GTGTCAATGCCTGAAGTGATACGCTCTGAGTGTCAATGCCTGAAGTGATACGCTCTGAGTGTCAATGCCTGAAGTGATACGCTCTGA | Bhatia et al.^3^ |
| N3 x 1 | CTTTGTTATAGGCTAAGCTCTCAGTAATCCCAAACAAAAGAGT | Inoue et al.^4^ |
| N3 x 3 | CTTTGTTATAGGCTAAGCTCTCAGTAATCCCAAACAAAAGAGTCTTTGTTATAGGCTAAGCTCTCAGTAATCCCAAACAAAAGAGTCTTTGTTATAGGCTAAGCTCTCAGTAATCCCAAACAAAAGAGT | Inoue et al.^4^ |
| DC5 x 1 | AAATATTCATTGTTGTTGCTCACCTACCATGGATCC | Hayashi et al.^5^ |
| DC5 x 3 | AAATATTCATTGTTGTTGCTCACCTACCATGGATCCAAATATTCATTGTTGTTGCTCACCTACCATGGATCCAAATATTCATTGTTGTTGCTCACCTACCATGGATCC | Hayashi et al.^5^ |
| DC5CON x 3 | AAATATTCATTGTTGATGTTCACGCATCATGGATCCAAATATTCATTGTTGATGTTCACGCATCATGGATCCAAATATTCATTGTTGATGTTCACGCATCATGGATCC | Narasimhan et al.^6^ |
| ACACA | AGGTTTGTATTCATTCTTTTCAGCTTGCTTGGATTTAGGTTTGTATTCATTCTTTTCAGCTTGCTTGGATTTAGGTTTGTATTCATTCTTTTCAGCTTGCTTGGATTT | Narasimhan et al.^6^ |
| P6CON x 3 | TTCAGGAAAAATTTTCACGCTTGAGTTCACAGCTCGAGTTTCAGGAAAAATTTTCACGCTTGAGTTCACAGCTCGAGTTTCAGGAAAAATTTTCACGCTTGAGTTCACAGCTCGAGT | Epstein et al.^7^ |
| CRY x 3 | ATTTCACGCATGAGTGCACATTTCACGCATGAGTGCACATTTCACGCATGAGTGCAC | Xu et al.^8^ |
| TFAP2A | GGAAGAACTGCCTGTTCACAATTTTCACCATACTGACCATTCGTTTTAAACGATTTTTTTCTTTTCTTTTCTGTGCTCTGACAGTCTGTTTGACTCAGATGCAGTTGTTTTCAAATAACATGAACGGAGCCTAACTGTACATGTCCTTGGCATAATTGCAATGCACAGTGCAGCAGCAGGTGAAGCGAGGCATTGTCTGTCAAGACTGTTTTTGTTGGCTGCTTCTCAGATTGTGTGTGGGATTCTAACCTCGGGCACAA | Coutinho et al.^8^ |
| FOXP2 | TGCGACTGGTCATTTGTGTTTAGGCCGCATAACATGTGGCTGTATGGAATTTACAGTGCGCCTCCAGTAGTAACTCCATTTCCCATCGGCCTCTCCAGAGACCAGGCGAAATGTCAAGCCGTGGCTTTGAAAGAGAGATCGAGCGAGGAGGGGGAATAGAGAGAGAGAGAGAAAGGGACAGGGTGTTATATCTGCTGGTAACAAACACTTATGAACATTACTCATGCTTGGGTGACCAGAGAC | Coutinho et al.^8^ |
| BLX x 3 | CAGTCAAGCGTACAGTCAAGCGTACAGTCAAGCGTA | Unpublished Data, FitzPatrick Lab |
| P6 SELEX x 3 | TGTGTTCACTCAAGCGGAAATGTGTTCACTCAAGCGGAAATGTGTTCACTCAAGCGGAAA | Unpublished Data, FitzPatrick Lab |
| FIMO x 3 | CAGTCAGGCGTGCAGTCAGGCGTGCAGTCAGGCGTG | Unpublished Data, FitzPatrick Lab |
| XL | CCTCCTTGAAACTGAAATTTTAGCATGTGATTAATTAACTTGTAATATTCTAATCAAGCTTATAAAAGAGCACTGTTGGGCGTGAGTGGAGGCGCCGGAAAAAAGCATCGAAAAAACTTAGAAAA | Redden et al.^10^ |
| Position 1 Mutagenesis primer | CTTGGCACAGCCGCCATGNNSAACAGCCACAGTGGAGTAAAC | This Study |
| Barcode_Amp_F | AATGATACGGCGACCACCGAGATCTACACNNNNNNACACTCTTTCCCTACACGACGCTCTTCCGATCTAGCGACAAGGAGGGCTGAGGACCGGTAGT | This Study |
| Barcode_Amp_R | CAAGCAGAAGACGGCATACGAGATNNNNNNGTGACTGGAGTTCAGACGTGTGCTCTTCCGATCTCTCGAGGCGCGCCATACTCA | This Study |
| Custom_Rd1_Seq | AGCGACAAGGAGGGCTGAGGACCGGTAGT | This Study |

##### Table S1: Oligonucleotide sequences

Rows 1 to 16 correspond to the bait sequences used to screen for PAX6-mediated antibiotic resistance, using either single (x1) or triple tandem repeats (x3). Row 17 represents a synthetic minimal yeast promoter. Rows 18 shows an example of a mutagenic oligonucleotide used to introduce all possible amino acids at position 2 in the PAX6 paired domain using a degenerate NNS sequence. Similar primers shifted at 3 nucleotide increments were used to cover all codons intended for mutagenesis in PAX6. Rows 19 to 20 depict the primers used to simultaneously amplify barcode DNA and append Illumina-compatible adapters. The 6N sequences correspond to unique dual indices used in demultiplexing during next-generation sequencing. Row 21 is the custom read 1 sequencing primer used in barcode sequence. N, any nucleotide; S, guanine or cytosine.

|  | **AUROC** | **n_Pathogenic** | **n_Benign** | **optimal** | **path_cor** | **beni_cor** | **path_inc** | **beni_inc** | **accuracy** | **AUBPRC** | **ROC_score** | **BPR_score** |
| --- | --- | --- | --- | --- | --- | --- | --- | --- | --- | --- | --- | --- |
| DMS_BLX_+Geneticin_ABS | 0.954 | 89 | 31 | 0.610 | 81 | 29 | 8 | 2 | 0.917 | 0.950 | 1.000 | 1.000 |
| MetaRNN | 0.932 | 91 | 31 | 0.978 | 77 | 29 | 14 | 2 | 0.869 | 0.923 | 0.966 | 0.948 |
| MutPred | 0.929 | 91 | 29 | 0.862 | 81 | 25 | 10 | 4 | 0.883 | 0.921 | 0.983 | 0.966 |
| DMS_BLX_-Geneticin_ABS | 0.926 | 89 | 31 | 1.014 | 81 | 27 | 8 | 4 | 0.900 | 0.922 | 0.947 | 0.930 |
| DMS_LE9_-Geneticin_ABS | 0.922 | 89 | 31 | 1.028 | 76 | 28 | 13 | 3 | 0.867 | 0.926 | 0.930 | 0.982 |
| DMS_LE9_+Geneticin_ABS | 0.913 | 89 | 31 | 1.443 | 77 | 29 | 12 | 2 | 0.883 | 0.903 | 0.912 | 0.860 |
| MutationAssessor | 0.901 | 91 | 31 | 3.070 | 82 | 24 | 9 | 7 | 0.869 | 0.902 | 0.897 | 0.897 |
| DMS_BLX_-Geneticin | 0.898 | 89 | 31 | 1.014 | 77 | 28 | 12 | 3 | 0.875 | 0.904 | 0.877 | 0.877 |
| CONDEL | 0.895 | 91 | 31 | 0.694 | 67 | 28 | 24 | 3 | 0.779 | 0.886 | 0.862 | 0.810 |
| VESPAl | 0.894 | 91 | 31 | 0.492 | 83 | 23 | 8 | 8 | 0.869 | 0.890 | 0.845 | 0.828 |
| DMS_LE9_-Geneticin | 0.892 | 89 | 31 | 0.977 | 73 | 28 | 16 | 3 | 0.842 | 0.903 | 0.807 | 0.842 |
| DMS_BLX_+Geneticin | 0.891 | 89 | 31 | -0.610 | 78 | 29 | 11 | 2 | 0.892 | 0.908 | 0.789 | 0.912 |
| MetaLR | 0.889 | 91 | 31 | 0.989 | 66 | 29 | 25 | 2 | 0.779 | 0.856 | 0.793 | 0.690 |
| ESM-1v | 0.886 | 91 | 31 | -12.026 | 86 | 23 | 5 | 8 | 0.893 | 0.852 | 0.759 | 0.672 |
| ClinPred | 0.886 | 91 | 31 | 0.991 | 84 | 23 | 7 | 8 | 0.877 | 0.823 | 0.741 | 0.621 |
| DMS_LE9_+Geneticin | 0.885 | 89 | 31 | -1.443 | 76 | 29 | 13 | 2 | 0.875 | 0.885 | 0.772 | 0.789 |
| CPT | 0.883 | 91 | 31 | 0.735 | 78 | 24 | 13 | 7 | 0.836 | 0.869 | 0.707 | 0.759 |
| DEOGEN2 | 0.878 | 91 | 31 | 0.969 | 80 | 24 | 11 | 7 | 0.852 | 0.857 | 0.707 | 0.707 |
| BayesDel | 0.876 | 91 | 31 | 0.479 | 74 | 27 | 17 | 4 | 0.828 | 0.803 | 0.690 | 0.586 |
| DeepSAV | 0.875 | 91 | 31 | 0.827 | 76 | 25 | 15 | 6 | 0.828 | 0.869 | 0.672 | 0.776 |
| M-CAP | 0.868 | 91 | 31 | 0.839 | 78 | 23 | 13 | 8 | 0.828 | 0.859 | 0.655 | 0.724 |
| SIFT4G | 0.868 | 37 | 23 | 0.025 | 36 | 19 | 1 | 4 | 0.917 | 0.795 | 0.561 | 0.439 |
| VARITY_R | 0.866 | 91 | 31 | 0.928 | 77 | 24 | 14 | 7 | 0.828 | 0.847 | 0.638 | 0.655 |
| VARITY_ER | 0.865 | 91 | 31 | 0.793 | 85 | 19 | 6 | 12 | 0.852 | 0.860 | 0.621 | 0.741 |
| REVEL | 0.858 | 91 | 31 | 0.908 | 82 | 23 | 9 | 8 | 0.861 | 0.825 | 0.603 | 0.638 |
| FATHMM | 0.846 | 91 | 31 | -6.070 | 72 | 25 | 19 | 6 | 0.795 | 0.822 | 0.586 | 0.603 |
| SIFT | 0.825 | 10 | 16 | 0.030 | 8 | 12 | 2 | 4 | 0.769 | 0.769 | 0.433 | 0.356 |
| EVE | 0.813 | 91 | 31 | 0.935 | 71 | 25 | 20 | 6 | 0.787 | 0.746 | 0.552 | 0.466 |
| DeepSequence | 0.812 | 91 | 31 | -7.092 | 88 | 19 | 3 | 12 | 0.877 | 0.750 | 0.534 | 0.466 |
| Polyphen2_HumVar | 0.802 | 91 | 30 | 0.999 | 54 | 26 | 37 | 4 | 0.661 | 0.762 | 0.500 | 0.534 |
| VEST4 | 0.796 | 91 | 31 | 0.887 | 61 | 27 | 30 | 4 | 0.721 | 0.742 | 0.466 | 0.448 |
| SuSPect | 0.794 | 91 | 31 | 60.000 | 70 | 24 | 21 | 7 | 0.770 | 0.766 | 0.448 | 0.517 |
| fathmm-XF | 0.782 | 91 | 31 | 0.964 | 78 | 22 | 13 | 9 | 0.820 | 0.758 | 0.466 | 0.517 |
| Eigen | 0.782 | 91 | 31 | 6.180 | 87 | 18 | 4 | 13 | 0.861 | 0.704 | 0.414 | 0.362 |
| MVP | 0.775 | 91 | 31 | 0.971 | 86 | 16 | 5 | 15 | 0.836 | 0.681 | 0.414 | 0.345 |
| Polyphen2_HumDiv | 0.763 | 91 | 27 | 0.996 | 78 | 16 | 13 | 11 | 0.797 | 0.690 | 0.397 | 0.345 |
| SNAP2 | 0.762 | 91 | 31 | 67.000 | 58 | 26 | 33 | 5 | 0.689 | 0.765 | 0.397 | 0.517 |
| CADD | 0.747 | 91 | 31 | 25.000 | 88 | 13 | 3 | 18 | 0.828 | 0.712 | 0.362 | 0.379 |
| PonP2 | 0.746 | 91 | 31 | 0.841 | 69 | 21 | 22 | 10 | 0.738 | 0.716 | 0.328 | 0.379 |
| fathmm-MKL | 0.743 | 91 | 31 | 0.956 | 69 | 21 | 22 | 10 | 0.738 | 0.724 | 0.310 | 0.431 |
| mutationTCN | 0.742 | 91 | 20 | -11.410 | 60 | 15 | 31 | 5 | 0.676 | 0.724 | 0.446 | 0.482 |
| BLOSUM62 | 0.728 | 79 | 28 | -1.000 | 66 | 15 | 13 | 13 | 0.757 | 0.660 | 0.298 | 0.246 |
| PROVEAN | 0.724 | 91 | 30 | -4.010 | 74 | 20 | 17 | 10 | 0.777 | 0.631 | 0.293 | 0.259 |
| LASSIE | 0.710 | 64 | 29 | 0.001 | 48 | 20 | 16 | 9 | 0.731 | 0.721 | 0.246 | 0.298 |
| Grantham | 0.696 | 91 | 31 | 60.000 | 72 | 17 | 19 | 14 | 0.730 | 0.621 | 0.259 | 0.172 |
| PonPS | 0.691 | 91 | 31 | 0.410 | 74 | 17 | 17 | 14 | 0.746 | 0.631 | 0.241 | 0.241 |
| phyloP | 0.683 | 91 | 31 | 7.189 | 73 | 20 | 18 | 11 | 0.762 | 0.621 | 0.224 | 0.207 |
| NetDiseaseSNP | 0.655 | 91 | 31 | 0.440 | 75 | 15 | 16 | 16 | 0.738 | 0.622 | 0.190 | 0.207 |
| sequence_unet | 0.639 | 91 | 31 | 0.205 | 69 | 16 | 22 | 15 | 0.697 | 0.666 | 0.172 | 0.276 |
| fitCons | 0.633 | 91 | 31 | 0.598 | 83 | 11 | 8 | 20 | 0.770 | 0.578 | 0.155 | 0.155 |
| DANN | 0.624 | 91 | 31 | 0.996 | 69 | 17 | 22 | 14 | 0.705 | 0.551 | 0.147 | 0.121 |
| MetaSVM | 0.616 | 91 | 31 | 1.001 | 65 | 20 | 26 | 11 | 0.697 | 0.507 | 0.138 | 0.017 |
| GenoCanyon | 0.610 | 91 | 31 | 1.000 | 83 | 10 | 8 | 21 | 0.762 | 0.568 | 0.103 | 0.138 |
| MutationTaster | 0.581 | 91 | 31 | 1.000 | 88 | 6 | 3 | 25 | 0.770 | 0.545 | 0.086 | 0.103 |
| SiPhy | 0.540 | 91 | 31 | 13.716 | 80 | 10 | 11 | 21 | 0.738 | 0.513 | 0.069 | 0.086 |
| PrimateAI | 0.538 | 91 | 31 | 0.876 | 68 | 14 | 23 | 17 | 0.672 | 0.510 | 0.052 | 0.034 |
| GERP++ | 0.524 | 91 | 31 | 3.550 | 91 | 6 | 0 | 25 | 0.795 | 0.511 | 0.034 | 0.069 |
| phastCons | 0.516 | 91 | 31 | 1.000 | 91 | 1 | 0 | 30 | 0.754 | 0.508 | 0.017 | 0.017 |
| LRT | 0.450 | 51 | 19 | 0.000 | 10 | 18 | 41 | 1 | 0.400 | 0.525 | 0.000 | 0.054 |

##### Table S2: Summary of variant classification by VEPs and DMS

Estimation of the ability of DMS assay data and VEPs to accurately classify pathogenic and benign PAX6 variants are shown as a measure of the area under the receiver operating characteristic curve (ROC AUC). Area under the precision-recall curve (AUBPRC) was also calculated. Normalised to the best performer, these are summarised as ROC_score and BPR_score, respectively. For accuracy and optimal thresholds determinations, see methods.
